# Large-scale analysis of SARS-CoV-2 synonymous mutations reveals the adaptation to the human codon usage during the virus evolution

**DOI:** 10.1101/2021.04.23.441151

**Authors:** Daniele Ramazzotti, Fabrizio Angaroni, Davide Maspero, Mario Mauri, Deborah D’Aliberti, Diletta Fontana, Marco Antoniotti, Elena Maria Elli, Alex Graudenzi, Rocco Piazza

**Author notes:** Equally contributing authors (Daniele Ramazzotti, Fabrizio Angaroni). Co-senior authors (Elena Maria Elli, Alex Graudenzi, Rocco Piazza).

## Abstract

Many large national and transnational studies have been dedicated to the analysis of SARS-CoV-2 genome, most of which focused on missense and nonsense mutations. However, approximately 30% of the SARS-CoV-2 variants are synonymous, therefore changing the target codon without affecting the corresponding protein sequence.

By performing a large-scale analysis of sequencing data generated from almost 400,000 SARS-CoV-2 samples, we show that silent mutations increasing the similarity of viral codons to the human ones tend to fixate in the viral genome over-time. This indicates that SARS-CoV-2 codon usage is adapting to the human host, likely improving its effectiveness in using the human aminoacyl-tRNA set through the accumulation of deceitfully neutral silent mutations.

**One-Sentence Summary:** Synonymous SARS-CoV-2 mutations related to the activity of different mutational processes may positively impact viral evolution by increasing its adaptation to human codon usage.

## Main Text

The COVID-19 pandemic caused by the Severe Acute Respiratory Syndrome Coronavirus 2 (SARS-CoV-2) has currently affected approximately 433 million people, with at least 5.9 million COVID-related deaths (1) (updated on March 1st, 2022), hence posing a major health threat on a global scale.

This led to a surge of active research aimed at studying SARS-CoV-2 biology: many large national and transnational efforts were dedicated to the analysis of SARS-CoV-2 genomes, with more than 1 million sequencing datasets currently available. These analyses succeeded in reconstructing the phylogeny of the virus and, at least in part, to implement effective lineage tracking strategies (2, 3, 4, 5, 6, 7). Notably, only a relatively small subset of the nonsynonymous mutations detected so far, with a few relevant exceptions such as the D614G Spike mutation (8), and the mutations associated with lineages B.1.1.7, B1.1.28, P1, B.1.617.2 and B.1.1.529 (9, 10, 11, 12) to name the major ones, were unequivocally shown to play a functional role. This evidence led to the conclusion that the majority of the reported SARS-CoV-2 sequence changes are neutral or detrimental, therefore having limited or no impact on the pathogenicity of COVID-19 pandemic (13).

To the best of our knowledge, most of the studies performed so far have focused on missense and nonsense mutations, i.e., mutations that either cause a change in the sequence of the target protein or generate a premature termination codon. However, a large fraction of the SARS-CoV-2 mutations (approximately ~30% in our datasets) are synonymous, therefore changing the target codon without affecting the associated protein sequence on mRNA translation, as opposed to missense/nonsense variants.

This bulk set of mutations is usually discarded during variant analyses, as they are considered as functionally irrelevant. However, it is known from decades (14, 15, 16, 17, 18, 19) that silent mutations may have a profound impact on protein expression by, i.e., changing the aminoacyl-tRNA molecule responsible for translating the mutated codon into the corresponding amino acid, a well-known process coordinated by the ribosomal machinery. A silent substitution causing a switch to a more abundant aminoacyl-tRNA is prone to increase the overall protein translating efficiency, while the opposite may occur if the tRNA pairing with the mutated codon is less abundant than the original one (20, 21, 22, 23, 24). Therefore, we hypothesize that synonymous SARS-CoV-2 variants may play a functional role by adapting the sequence of the virus to the codon usage (CU) of the human host, thus improving its effectiveness in using the human aminoacyl-tRNA set, ultimately making the translation of viral particles more efficient (25).

Notice that SARS-CoV-2 variants might be classified into *minor variants*, i.e., detected with a low variant frequency (VF) within a host (i.e., VF < 90%) and *fixed variants,* i.e., detected with high variant frequency (i.e., VF > 90%). Variant frequency profiles can be derived via variant calling from raw sequencing data of viral samples, as proposed, i.e., in (5,6).

Owing to the on complex viral transmission dynamics involving bottlenecks and founder effects (5,6), mutations, originating in a small fraction of the virion pool, are fixed over-time only in presence of positive selection, otherwise they tend not to fixate in the population.

To investigate the impact of codon usage adaptation to the SARS-CoV-2 virus evolution, we analyzed a total of 390,899 viral samples and 3,178,178 silent viral mutations identified in almost 200 different studies (see Materials and Methods), showing that:

1. The viral adaptation to the human CU is showing an increasing trend over time during the pandemic.
2. Mutations with high codon adaptation are preferred and have a higher chance of fixating in the (intra-host) population.

To analyze a possible trend leading to the adaptation of the viral codons to the human CU during COVID-19 pandemic, we initially analyzed a subset of high-quality samples from North America (Dataset 1, see Materials and Methods). This dataset comprises a total of 213,737 COVID-19 cases, where viral SARS-CoV-2 RNA was sequenced using an Amplicon strategy with high coverage. After alignment of the raw viral data to the SARS-CoV-2-ANC reference genome (5) and variant calling (see Materials and Methods for further details), silent variants were annotated in terms of reference and mutated codon, variant frequency (VF) and sampling time (Supplementary Table 1), with ‘reference codon’ indicating the codon found in the SARS-CoV-2-ANC reference genome and ‘mutated codon’ the one generated by the presence of a single nucleotide variant. The relative codon usage (RCU) adaptation was calculated as the ratio between the human CU of the mutated codon and the human CU of the wild-type counterpart, with values > 1 indicating positive adaptation.

We analyzed the relative codon usage profile of mutated *vs* wild-type codons in the North America subgroup. Interestingly, we noticed that the APOBEC variants (C>T and G>A mutations) are both consistently associated with a negative value of log2-transformed RCU (see Figure 1 and Supplementary Figures 1 and 2, median value −0.30), i.e., leading to a reduction in the adaptation to the human codon usage. As previously shown by our team and by others (6, 26, 27, 28), a relevant subset of minor variants, linked to the C>T substitution type, are most likely generated by the activation of the human APOBEC machinery (despite other mutational processes cannot be excluded). APOBEC represents a family of cytidine deaminases playing a critical role in the intrinsic responses of the host to infections operated by a large set of viruses, among which retroviruses, herpesviruses, papillomaviruses, parvoviruses, hepatitis B virus and retrotransposons. APOBEC cytidine deaminases target the viral genome, eventually causing the functional inactivation of the pathogen. Therefore, APOBEC mutations globally represent a defense raised by the host cells to suppress viral infection (25).

**Figure 1.**
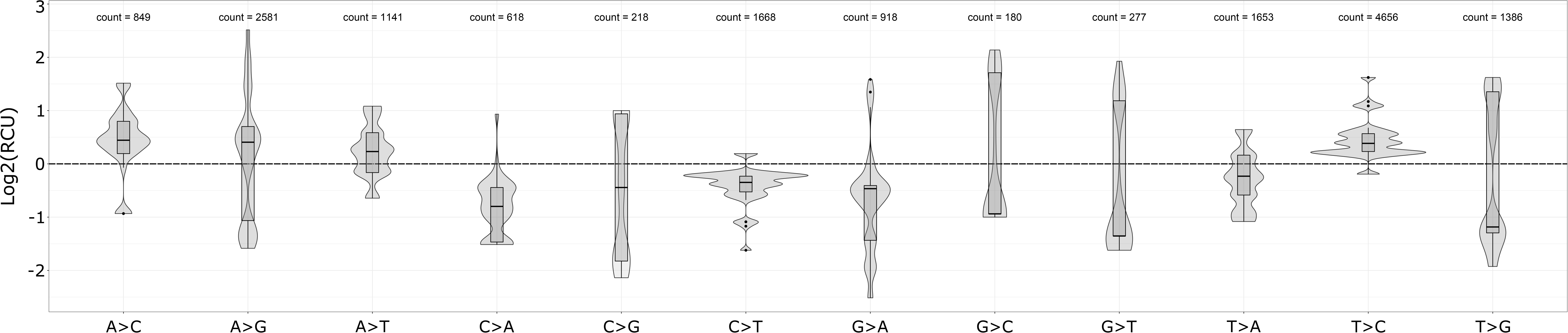
Empirical log2-transformed RCU distribution of all unique detected substitutions divided by substitution types. Violin and box plots representing the empirical distribution of the log2-transformed RCU values for each unique variant observed in Dataset 1; mutations observed in multiple samples are considered only once. The mutations were divided into 12 classes related to the possible nucleotide substitutions. For each variant type the overall number of all unique detected substitutions is reported.

**Figure 2.**
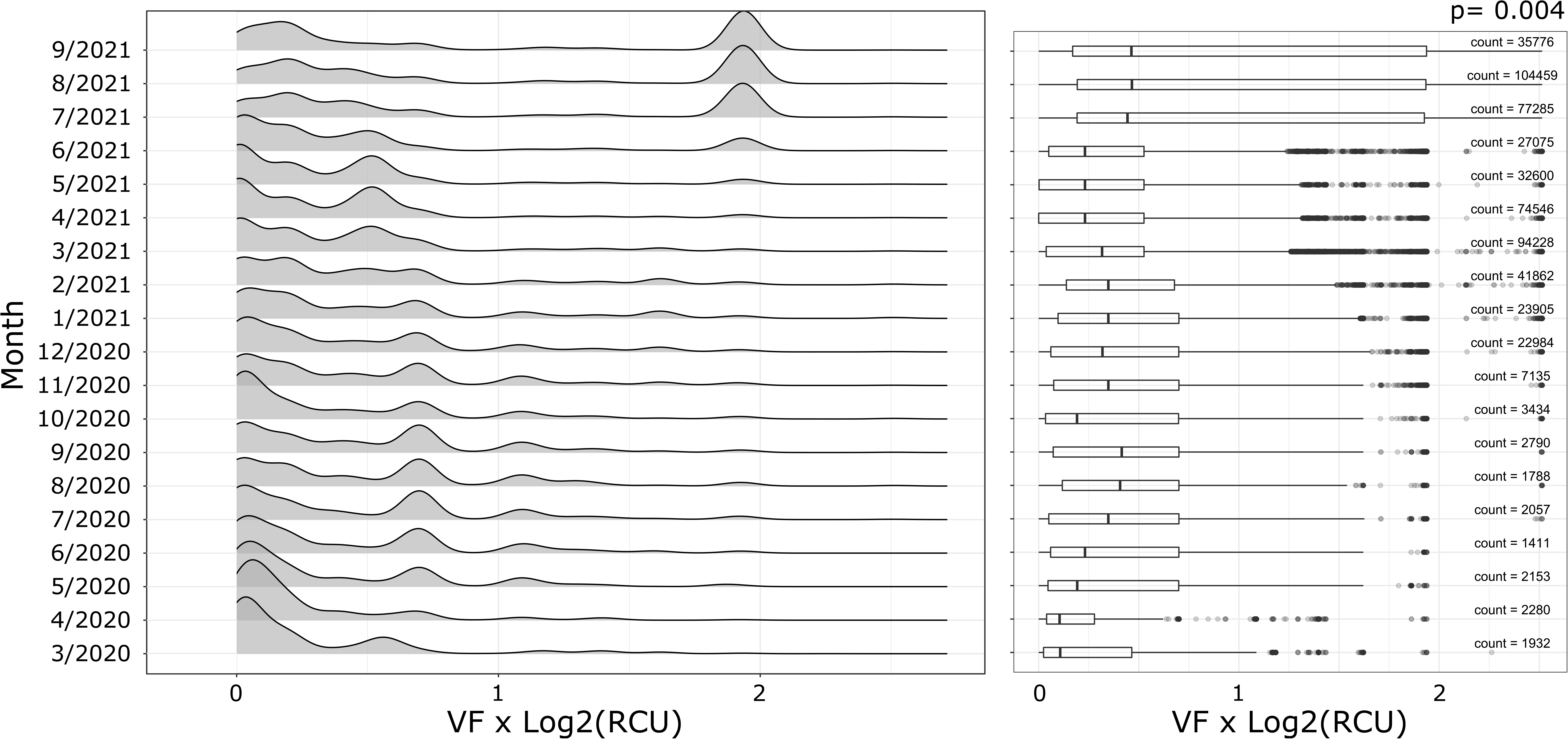
Temporal trend of the log2-transformed RCU for all the substitutions with positive codon adaptation with respect to the human CU. On the left, we report the distribution of the log2-transformed RCU multiplied by the VF of every mutation with a positive adaptation to the human CU, binned (per month) with respect to collection date for Dataset 1. On the right, the boxplots of the same distribution are shown and annotated with the number of observations for Dataset 1. The p-value reported on the top right of the Figure is computed performing a Mann-Kendall test for monotonic trend over the mean of the distribution for each month.

In Figure 1, we report the log2-transformed RCU for all the 12 possible mutation classes, i.e., all the possible mutations involving the four bases (6). As one can notice, different mutation classes may constitutively show different trends in terms of RCU. As an example, let us consider the variants likely linked to mutations generated by the RNA-specific adenosine deaminase ADAR (even if other mutational processes cannot be excluded) (6), which show an opposite behavior compared to the APOBEC ones. ADAR represent one of the other major observed sources of mutations in SARS-CoV-2 genomes and are associated to A>G and T>C mutations (6); such variants show a positive trend of adaptation to the human codon usage with a median value of +0.58 log2-transformed RCU (see Figure 1 and Supplementary Figures 1 and 2; APOBEC *vs* ADAR log2-transformed RCU t-test p-value <0.001), indicating that synonymous mutations related to different mutational processes may lead to very different effects in terms of CU.

A similar pattern was detected in two additional datasets: the first one (Dataset 2, Supplementary Figures 1 and 2) comprising 118,386 samples from the United Kingdom and the second one (Dataset 3, Supplementary Figures 1 and 2) including 58,776 samples from the rest of the world (see Materials and Methods for details). All the analyses corroborated our hypothesis, confirming in these two additional datasets all the results and conclusions discussed above.

Furthermore, these results do not change significantly when the CoCoPUTs human codon usage reference dataset from the Hive Lab (https://hive.biochemistry.gwu.edu/cuts/about; 29, 30) is considered for the analyses in place of the classical Kazusa codon usage reference (https://www.kazusa.or.jp/codon/cgi-bin/showcodon.cgi?species=9606; 31).

As different mutational processes may lead to different effects in terms of CU, we hypothesized that a RCU less fit to the human translational machinery could cause mutated SARS-CoV-2 genomes to reduce their translation efficiency, therefore undergoing purifying selection. As opposite, the variants leading to a better match with the human codon usage should undergo positive selection, overcome translational bottlenecks and outcompeting wild-type viruses in protein synthesis and ultimately improving the efficiency of viral packaging. Although a direct demonstration of this hypothesis is challenging, a greater RCU should correspond to an increased viral fitness, therefore potentially translating into a greater VF at the global level. To shed light on this phenomenon, we reasoned that the process of synonymous variants selection over time could be detected in very large time-series analyses. Accordingly, we binned all the silent variants based on their collection date in months intervals and we analyzed the trend of the VF for variants with positive or negative log2-transformed RCU, where positive values indicate increase in the similarity to the human CU and negative values the opposite. In particular, we first divided the mutations in two groups, i.e., the ones with a negative RCU (among which we have the majority of the APOBEC-associated mutations) and the ones with a positive RCU. We then grouped variants by collection date (months) and analyzed for each of the two groups the presence of any statistical trend over time of their RCU multiplied by VF, and finally log2-transformed. In line with the original hypothesis, silent variants in the positive CU group (Figure 2 and Supplementary Figures 3 to 7) showed an increase of the log2-transformed VF-weighted RCU over time, significant for all three datasets (one-sided Mann-Kendall test, p-value=0.004, 0.006, and 0.004 for Dataset 1, Dataset 2, and Dataset 3 respectively), suggesting that CU adaptation might play a significant role in the evolution of SARS-CoV-2 virus. As expected, no increase could be detected for the variants in the negative CU group. In conclusion, the results provided here point at the evidence that codon usage adaptation may play a role in the dynamics of minor mutations transitioning to fixed, in addition to functional selection.

Furthermore, to provide additional evidence to confirm that codon adaptation is improving when acquiring mutations over the course of the pandemic, we performed two additional analyses for all the 3 considered datasets:

1. With the aim of evaluating whether, in general, variants leading to better codon adaptation are preferred with respect to the ones not improving codon usage, we performed the following analysis. We considered the genome position of 100,000 randomly selected mutations (both synonymous and nonsynonymous) present in Dataset 1, Dataset 2 and Dataset 3 (e.g., position 150 for variant 150 C>T), tested all the alternative nucleotide substitutions for that position (i.e., 150 C>G and 150 C>A), and checked whether each of such substitutions would lead to the synthesis of the same protein generated by the original variant (i.e., 150 C>T). This led us to a set of variants where different mutations lead to the synthesis of the same protein. Within this set, we compared the distribution of the codon usage values obtained with the original variants to the one that would be achieved with the alternative substitutions by performing a standard t-test, which led to a very significant p-value (< 0.0001) for all the three datasets. When repeating the same analysis with the CoCoPUTs human codon usage, the results were confirmed.
2. We first split the timeline covered by the datasets in two subsequent time frames: (*i*) first half of the timeline (2020/01/01 to 2020/11/30 for Dataset 1; 2020/01/01 to 2020/10/31 for Dataset 2; 2020/01/01 to 2020/12/31 for Dataset 3), (*ii*) second half of the timeline (until 2021/09/30 for Dataset 1, 2021/07/31 for Dataset 2 and until 2021/11/30 for Dataset 3); we selected the variants displaying a VF > 0.90 for the first time in either one of the two time frames (i.e., fixed variants); we finally compared the number of such variants belonging to the APOBEC group. Specifically, we reported: for Dataset 1 41.45% of APOBEC variants in time frame (*i*) and 11.13 in time frame (*ii*); for Dataset 2 42.34% of APOBEC variants in time frame (*i*) and 15.17 in time frame (*ii*); for Dataset 3 38.71% of APOBEC variants in time frame (*i*) and 16.19 in time frame (*ii*). We finally performed a standard z-test to compare the proportions for each time frame and obtained a highly significant p-value (<0.0001) for all the three datasets, confirming that APOBEC variants (with negative codon adaptation, on average) show reduced trend of fixation in the population as the pandemic continues, hinting at the evidence that mutations showing negative codon usage have lower likelihood of transitioning from minor to fixed.

Taken globally, these results suggest that silent mutations may play a role in viral evolution, by increasing the adaptation of the viral genome to the human codon usage. To this end, it is tempting to speculate that the observed continuous adaptation of its codon usage over time may underlie an increase of the overall efficiency of viral protein production and packaging, with viral genomes with a better adaptation being able to generate more viral particles over time, therefore outperforming other, less adapted, viruses. It is widely accepted that improvements in codon usage adaptation, by removing translation bottlenecks, may contribute to the overall protein synthesis efficiency (32). Indeed, proteins differing only at silent sites can have several orders of magnitude of difference in their expression level (33). This finding has been widely exploited in both research and industry to adapt coding sequences to specific organisms to optimize protein synthesis. Recently, codon usage optimization was also proposed as an effective strategy to drive high levels of viral protein expression in human cells to stimulate a very robust immune response (34). Recent studies pointed also to more complex scenarios, where the use of different codons may allow to fine-tune the translation time of individual gene regions, to allow proper folding of complex domains as well as to maximize the occurrence of post-translational modifications (35); in this context a very fast translational process is not always an advantage, as in specific conditions pausing the translation machinery may be beneficial for the production of functional, properly folded proteins. Further analyses and very large datasets will be required to identify these specific events, if present.

As the number of available viral sequences is quickly increasing over time, it will be similarly interesting to isolate individual silent variants showing relevant changes in their VF over large time spans to study specific codon usage adaptation patterns in the context of individual viral genes/proteins. Notably, the effect of codon adaptation could be also directly assessed in terms of viral fitness. By using a reverse genetics approach, *in vitro* modified viral RNA genomes could be generated carrying progressively more human-adapted genomes. The functional effect of these modifications could be then assessed in terms of increased viral fitness or in competition assays. Similarly, the overall level of protein expression could be tested in target cells by using conventional approaches such as western blot and confocal microscopy.

In conclusion, global as well as time-series analyses of silent mutations indicate that codon usage adaptation may play a relevant role in the evolution of SARS-CoV-2, suggesting that the evolution of the viral genome has been intense. Further studies will be required to thoroughly dissect the overall impact and clinical relevance of these findings. Soon, it will be important to assess if specific viral genomes with a high codon usage adaptation also show an increased infectiousness and/or clinical aggressiveness, which, at the time being, is very difficult, owing to the limited availability of clinical data in publicly available SARS-CoV-2 sequencing studies. In this regard and given the profound impact of COVID-19 pandemic, every effort should be made to share clinical information together with sequencing data, which is unfortunately rarely done.

## Acknowledgments

We acknowledge that this research was partially supported by the Italian Ministry of University and Research (MIUR) - Department of Excellence project PREMIA (PREcision MedIcine Approach: bringing biomarker research to the clinic).

## Funding

Italian Ministry of University and Research (MIUR) - Department of Excellence project PREMIA (PREcision MedIcine Approach: bringing biomarker research to the clinic). This work was partially supported by a Bicocca 2020 Starting Grant to FA and DR.

## Author contributions

Conceptualization: RP
Methodology: RP, DR, AG
Investigation: DR, FA, DM
Visualization: FA, DM, DR
Funding acquisition: RP, AG, DR
Supervision: RP, AG, EME, DR
Writing – original draft: DR, FA, AG, RP
All authors read and approved the final manuscript.

## Competing interests

Authors declare that they have no competing interests.

## Data and materials availability

All data are available in the main text or the supplementary materials or can be downloaded from the original SARS-CoV-2 sequence repositories.

## Supplementary Materials

Materials and Methods

Figs. S1 to S7

Table S1

## Notes

https://github.com/danro9685/codon_usage

### Competing Interest Statement

The authors have declared no competing interest.

