## Supplementary Materials for "Large-scale analysis of SARS-CoV-2 synonymous mutations reveals the adaptation to the human codon usage during the virus evolution"

+ Equally contributing authors

† Co-senior authors

**This PDF file includes:**

Materials and Methods
Figs. S1 to S7
Table S1

### Materials and Methods

#### Datasets

**Dataset #1.** We analyzed a dataset obtained by considering data from 68 distinct NCBI BioProjects. For all samples, Illumina AMPLICON paired-end high-coverage sequencing data are provided; all patients were in the United States. Within this dataset, we considered for our analyses 213,737 high-quality samples having coverage ≥ 100 in at least 90% of the virus genome, collected between January 2020 and September 2021.
**Dataset #2.** We analyzed a dataset obtained by considering data from 8 distinct NCBI BioProjects. For all samples, Illumina AMPLICON paired-end high-coverage sequencing data are provided; all patients were in the United Kingdom. Within this dataset, we considered for our analyses 118,386 high-quality samples having coverage ≥ 100 in at least 90% of the virus genome, collected between January 2020 and July 2021.
**Dataset #3.** We analyzed a dataset obtained by considering data from 128 distinct NCBI BioProjects. For all samples, Illumina AMPLICON paired-end high-coverage sequencing data are provided; patients were from multiple locations around the world, except for the United States and the United Kingdom which were included in the previous datasets. Within this dataset, we considered for our analyses 58,776 high-quality samples having coverage ≥ 100 in at least 90% of the virus genome, collected between January 2020 and November 2021.

#### Variant calling

Variant calling was performed by employing the iVar (version 1.3.1) recommended pipeline to analyze SARS-CoV-2 ARTIC v3 amplicon sequencing data. We performed the following steps: 1) FASTQ files were mapped to the reference genome SARS-CoV-2-ANC with *bwa mem* (version 0.7.17-r1188). 2) Sorted BAM files were generated from bwa mem results using SAMtools (version 1.10). 3) ARTICv3 primer sequences were trimmed using the *ivar trim* command. 4) Trimmed sorted BAM files were built and indexed with SAMtools. 5) Mutation calling was performed from trimmed sorted BAM files using *ivar variants*. 6) Finally, *samtools depth* was used to extract coverage information from trimmed sorted BAM files.
Quality control was performed on the mutations obtained with *ivar variants*. We first selected (ultra) deep sequencing samples with a coverage of at least 100 reads in at least 90% of the viral genome. Then, we performed further filtering by selecting only mutations with variant frequency of at least 5%, where mutations were supported by at least 10 reads and with p-value resulting from the *ivar variants* algorithm less than 0.01. Finally, samples with more than 100 minor mutations (after filtering) were removed.

#### Software availability

The source code used to replicate all the analyses presented in this work is available at this link: https://github.com/danro9685/codon_usage.

**Figure S1. Empirical log2-transformed RCU distribution of all unique detected substitutions divided by substitution types for datasets 2 and 3**. Violin and box plots representing the empirical distribution of the log2-transformed RCU values for each unique variant observed in Dataset 2 (panel A) and Dataset 3 (panel B); mutations observed in multiple samples are considered only once. The mutations were divided into 12 classes related to the possible nucleotide substitutions. For each variant type the overall number of all unique detected substitutions is reported.

**Figure S2. Empirical log2-transformed RCU distribution of all detected substitutions divided by substitution types for the three datasets.** Violin and box plots representing the empirical distribution of the log2-transformed RCU values for all variants observed in Dataset 1 (panel A), Dataset 2 (panel B) and Dataset 3 (panel C); mutations observed in multiple samples are considered multiple times. The mutations were divided into 12 classes related to the possible nucleotide substitutions. For each variant type the overall number of all detected substitutions is reported.

**Figure S3. Temporal trend of the log2-transformed RCU for all the substitutions with negative codon adaptation with respect to the human CU for Dataset 1**. On the left, we report the distribution of the log2-transformed RCU multiplied by the VF of every mutation with a negative adaptation to the human CU, binned (per month) with respect to collection date for Dataset 1. On the right, the boxplots of the same distribution are shown and annotated with the number of observations for Dataset 1. The p-value reported on the top right of the Figure is computed performing a Mann-Kendall test for monotonic trend over the mean of the distribution for each month.

**Figure S4. Temporal trend of the log2-transformed RCU for all the substitutions with positive codon adaptation with respect to the human CU for Dataset 2.** On the left, we report the distribution of the log2-transformed RCU multiplied by the VF of every mutation with a positive adaptation to the human CU, binned (per month) with respect to collection date for Dataset 2. On the right, the boxplots of the same distribution are shown and annotated with the number of observations for Dataset 2. The p-value reported on the top right of the Figure is computed performing a Mann-Kendall test for monotonic trend over the mean of the distribution for each month.

**Figure S5. Temporal trend of the log2-transformed RCU for all the substitutions with negative codon adaptation with respect to the human CU for Dataset 2.** On the left, we report the distribution of the log2-transformed RCU multiplied by the VF of every mutation with a negative adaptation to the human CU, binned (per month) with respect to collection date for Dataset 2. On the right, the boxplots of the same distribution are shown and annotated with the number of observations for Dataset 2. The p-value reported on the top right of the Figure is computed performing a Mann-Kendall test for monotonic trend over the mean of the distribution for each month.

**Figure S6. Temporal trend of the log2-transformed RCU for all the substitutions with positive codon adaptation with respect to the human CU for Dataset 2.** On the left, we report the distribution of the log2-transformed RCU multiplied by the VF of every mutation with a positive adaptation to the human CU, binned (per month) with respect to collection date for Dataset 3. On the right, the boxplots of the same distribution are shown and annotated with the number of observations for Dataset 3. The p-value reported on the top right of the Figure is computed performing a Mann-Kendall test for monotonic trend over the mean of the distribution for each month.

**Figure S7. Temporal trend of the log2-transformed RCU for all the substitutions with negative codon adaptation with respect to the human CU for Dataset 2.** On the left, we report the distribution of the log2-transformed RCU multiplied by the VF of every mutation with a negative adaptation to the human CU, binned (per month) with respect to collection date for Dataset 3. On the right, the boxplots of the same distribution are shown and annotated with the number of observations for Dataset 3. The p-value reported on the top right of the Figure is computed performing a Mann-Kendall test for monotonic trend over the mean of the distribution for each month.

**Table S1. Synonymous variants data with associated sampling date and codon usage information.**
