## Supplementary figures and images for "Large-scale analysis of SARS-CoV-2 synonymous mutations reveals the adaptation to the human codon usage during the virus evolution"

### Figure S1

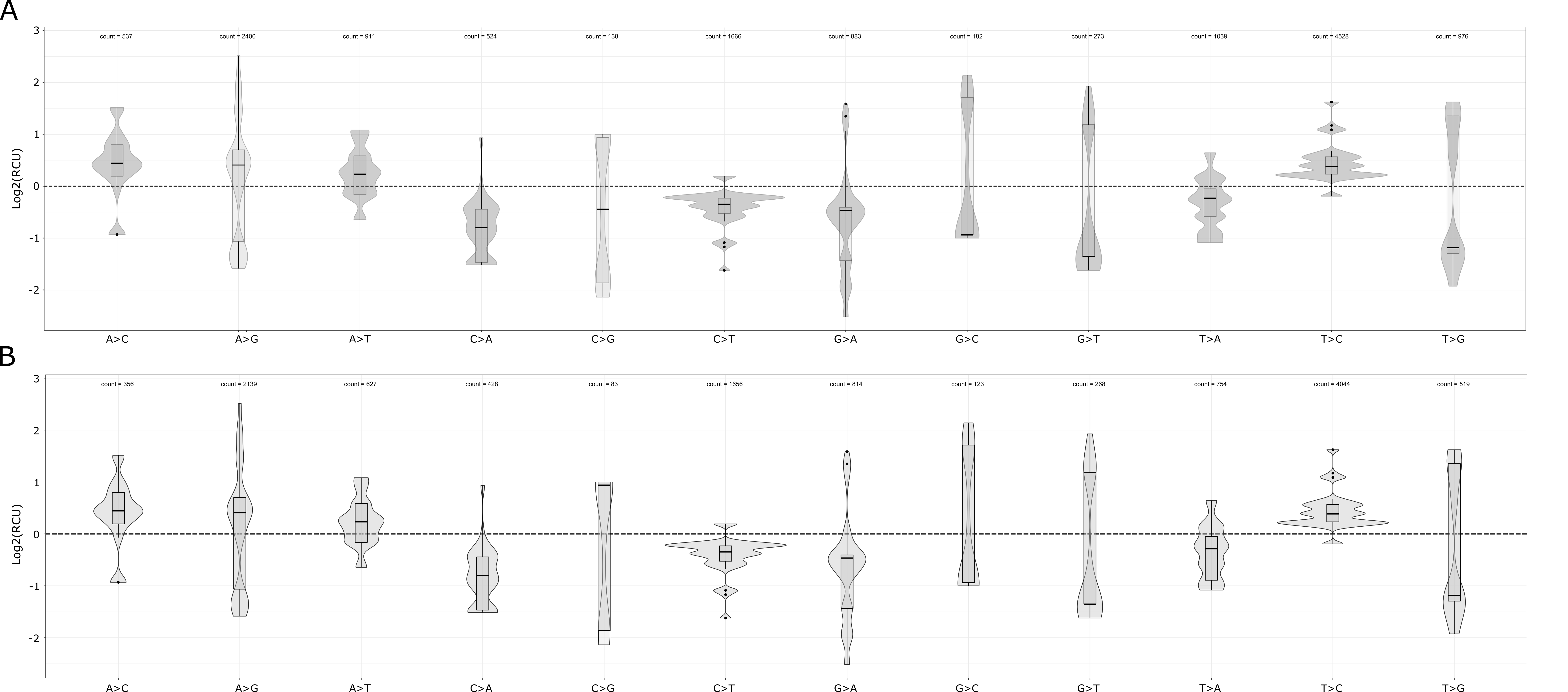

### Figure S2

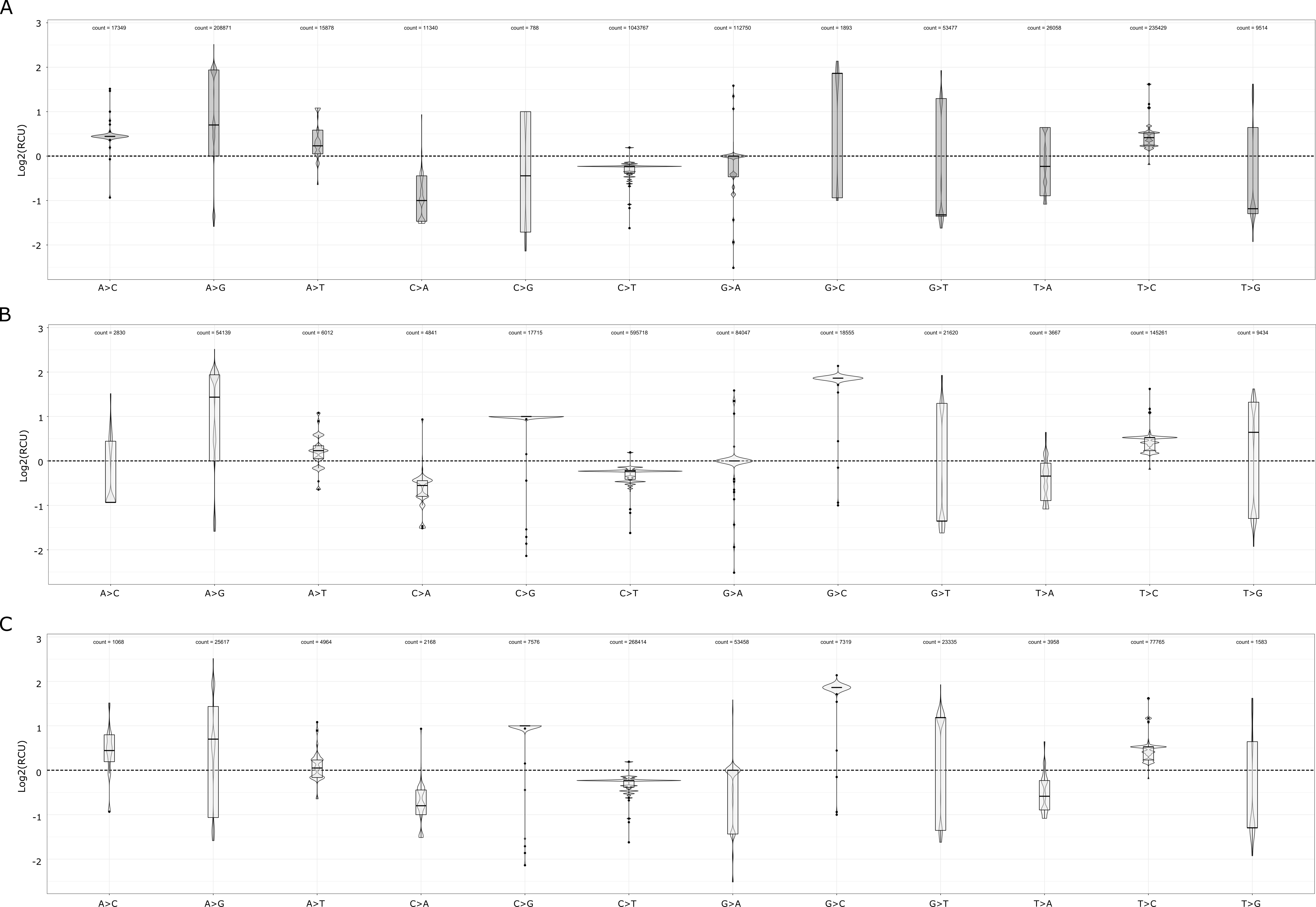

### Figure S3

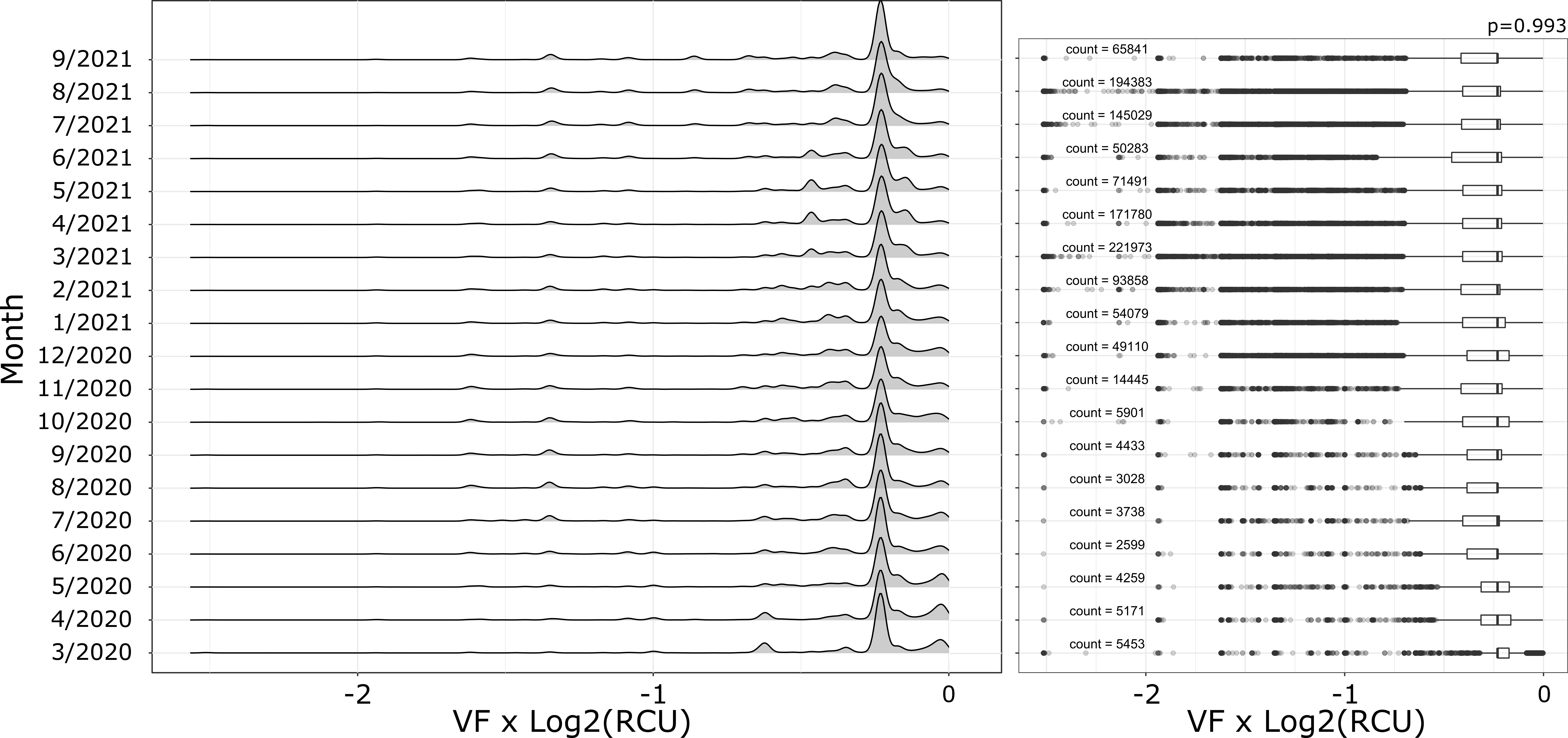

### Figure S4

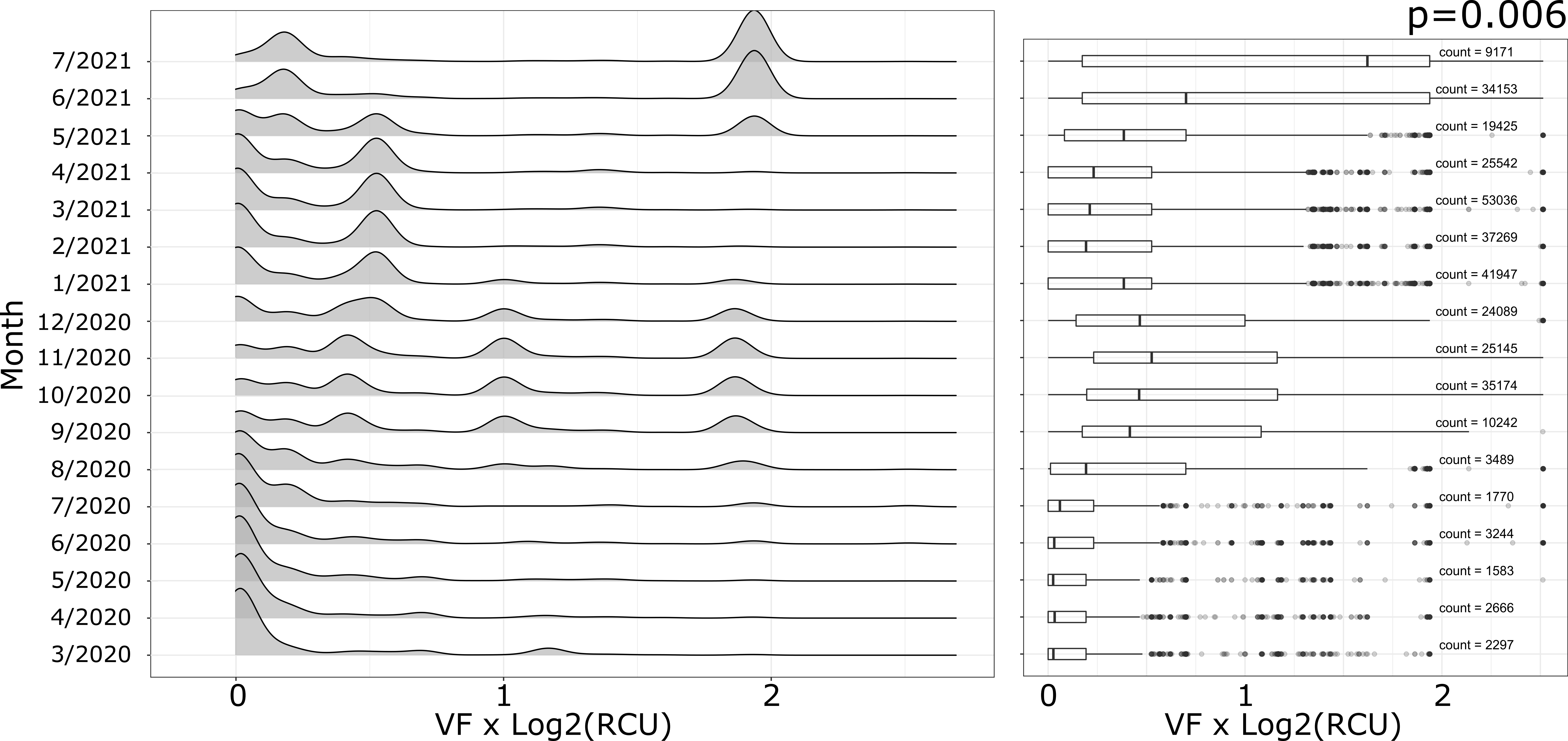

### Figure S5

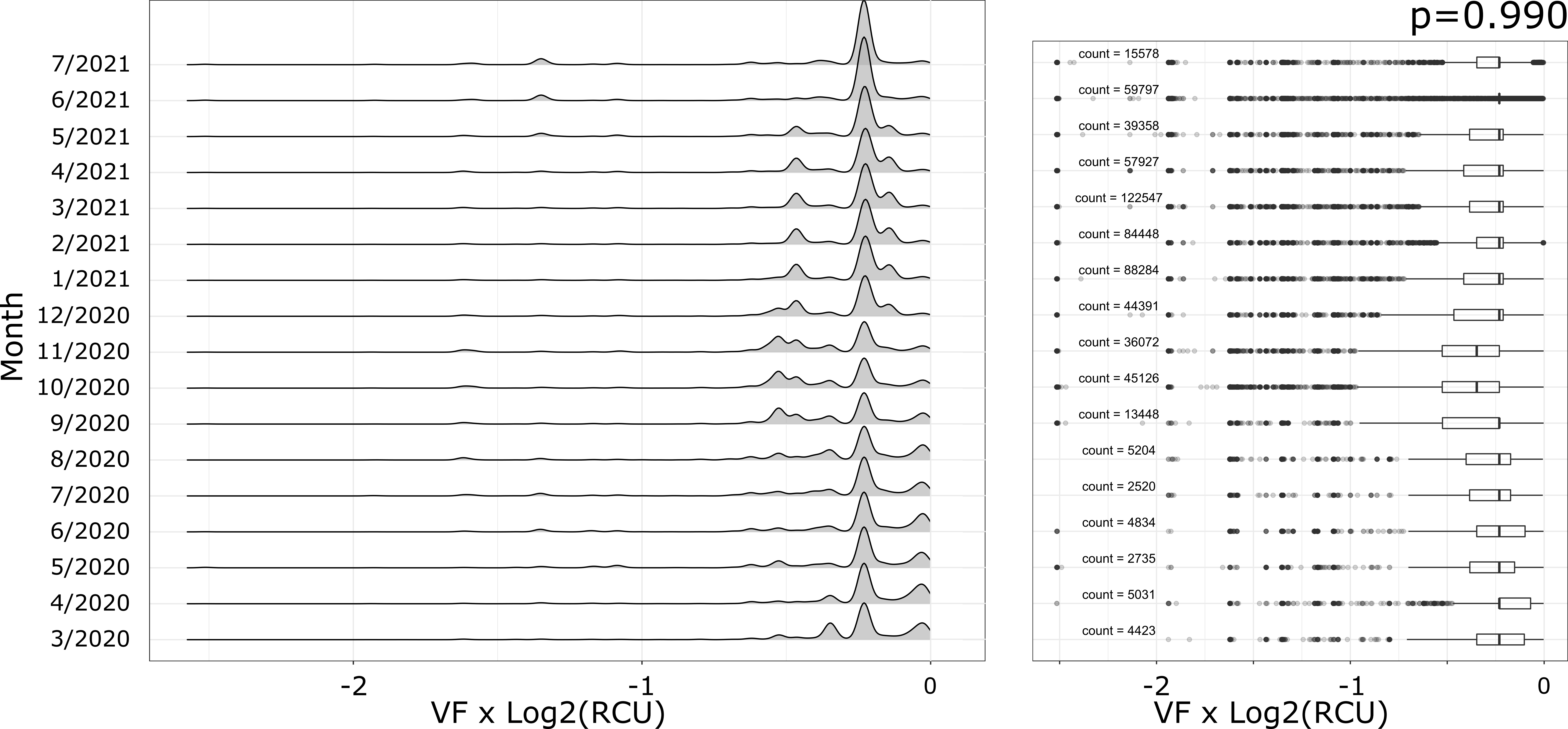

### Figure S6

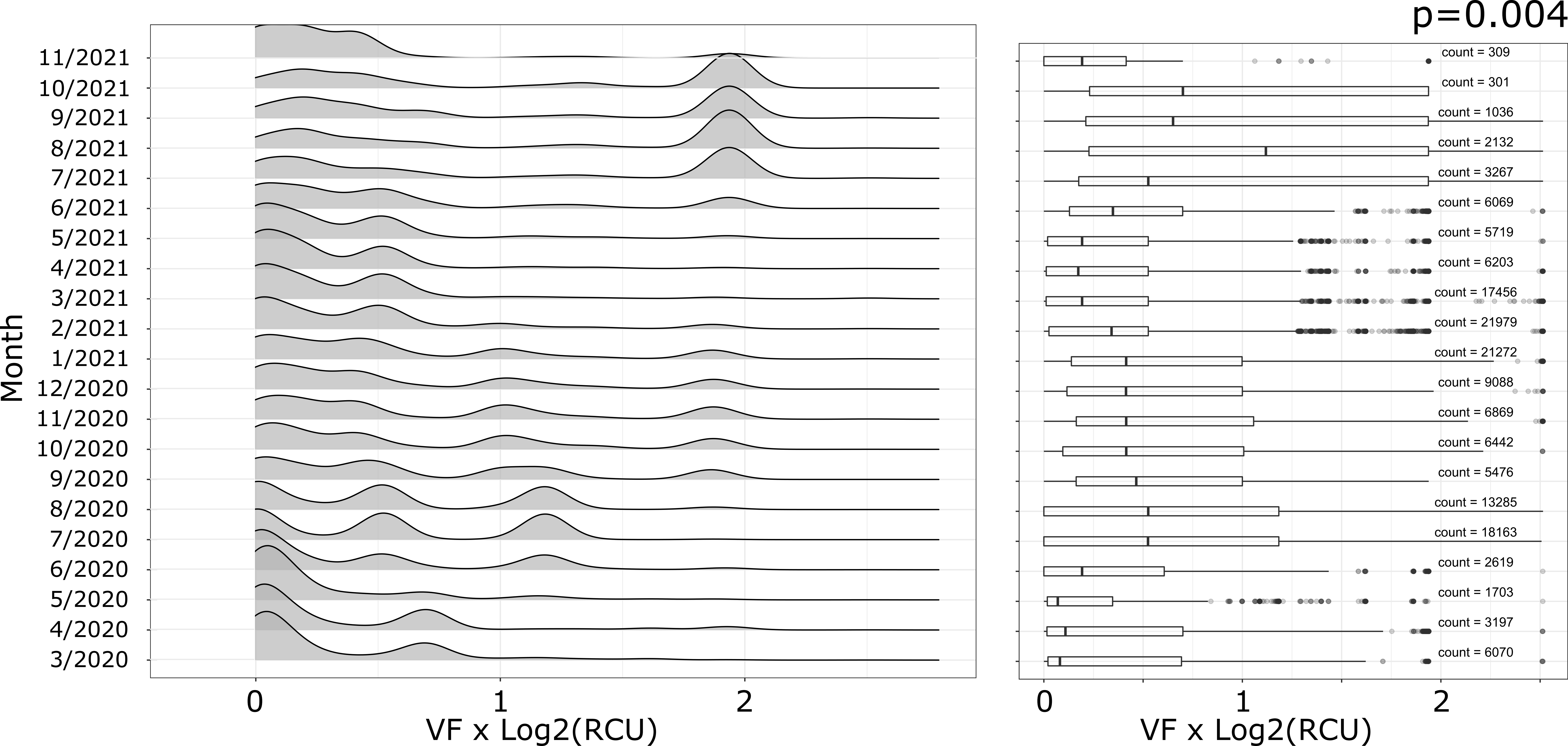

### Figure S7

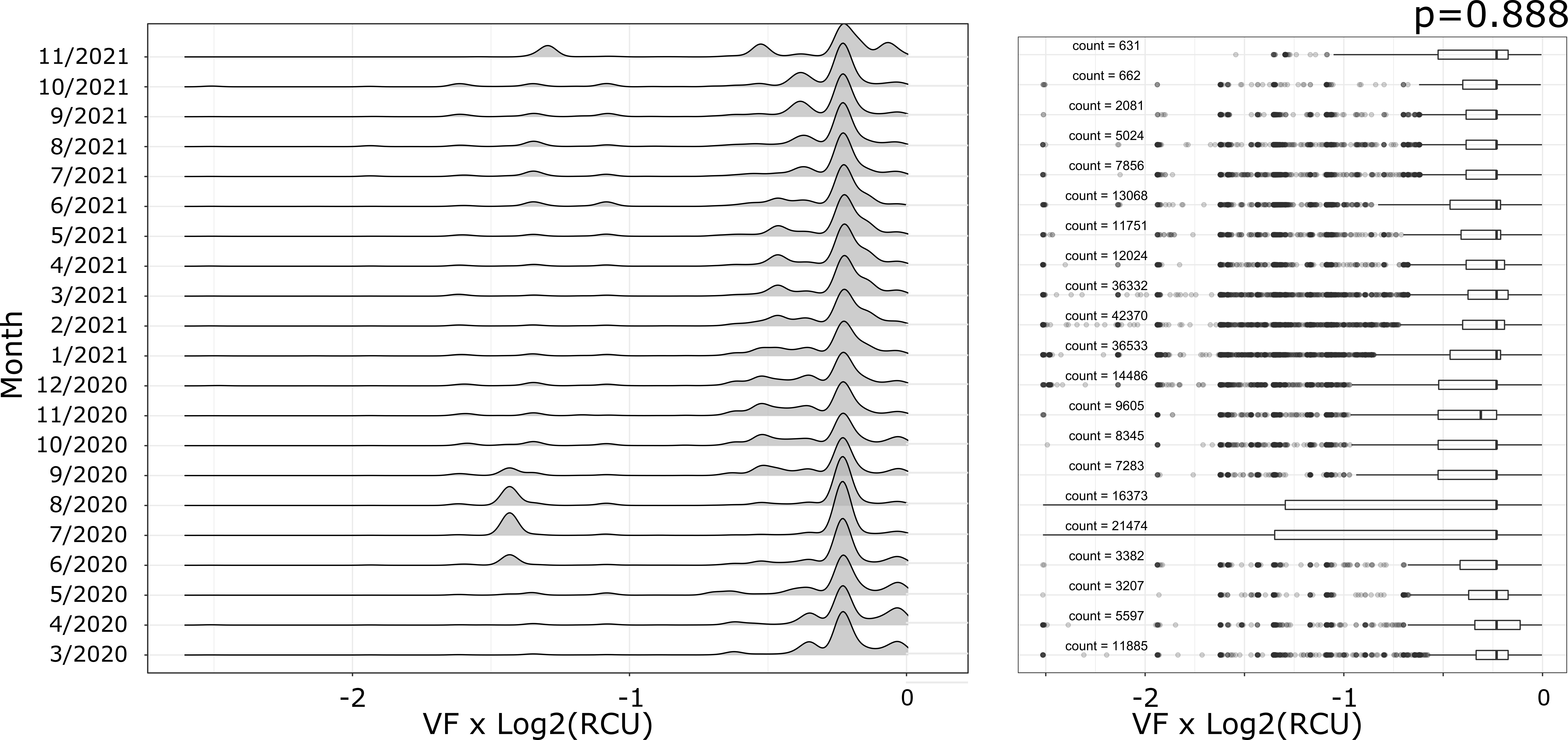
